# Go/z-biased coupling profile of the dopamine D3 receptor

**DOI:** 10.1101/2025.08.08.668522

**Authors:** Lucrezia Zanetti, Luca Franchini, Shirsha Saha, Yini Liao, Brian S. Muntean, Cesare Orlandi

## Abstract

Dopamine receptors are G protein coupled receptors (GPCRs) that serve as key targets for FDA-approved drugs used to treat various neuropsychiatric disorders. Notably, ∼11% of all marketed GPCR-targeting drugs act on dopamine receptors. Five GPCRs mediate the effects of endogenous dopamine and compounds used to treat Parkinson’s disease, schizophrenia, and other conditions. However, on-target side effects associated with these medications highlight the need to analyze dopamine receptor signaling to design safer, more effective therapeutics.

We characterized the G protein coupling of dopamine D2-like receptors and observed the striking inability of D3R to engage with G_i_ proteins while effectively activating G_o_ and G_z_ subtypes. Applying orthogonal cell-based assays that utilize wild-type G proteins both in parental and ΔGα_i/o/z_ cells, we conclusively established that D3R does not activate G_i_ proteins. Further analysis of Gα_i2_:Gα_oA_ and D2R:D3R chimeras revealed that this selective inability is driven by molecular determinants located within the α5 helix of Gα_i_ and the intracellular loop 2 (ICL2) of D3R. Guided by cryo-EM structures, we modeled the interface between these regions to better understand the structural basis of this selectivity. Finally, we treated hippocampal neurons in acute brain slices with selective agonists for D2R and D3R and observed marked differences in their ability to regulate endogenous adenylyl cyclase to produce cAMP, highlighting the neurophysiological significance of our findings.

## INTRODUCTION

Dopamine is a critical catecholaminergic neurotransmitter that regulates a broad range of functions in both the central and the peripheral nervous systems^1–3^. Dopamine actions are mediated by the activation of five G protein-coupled receptors (GPCRs) that are classified into two families based on their structural and signaling properties: D1-like receptors (D1R, D5R) and D2-like receptors (D2R, D3R, D4R). D1-like receptors generally couple to heterotrimeric G_s_ proteins that stimulate adenylyl cyclase (AC) to produce cAMP, whereas D2-like receptors predominantly couple to inhibitory G_i/o/z_ proteins, resulting in AC inhibition and activation of additional downstream signaling pathways, including those mediated by β-arrestin recruitment^3–11^. A key emerging concept in signal transduction is that a single receptor can adopt multiple conformations, each selectively engaging different intracellular transducers and triggering distinct cellular responses^12–15^. This signaling bias includes preference not only between G protein and β-arrestin pathways but also among different G protein subtypes^16–19^. Dysregulation of D2-like receptor signaling has been implicated in a wide range of neurological and psychiatric disorders, including Parkinson’s disease (PD), Huntington’s disease (HD), schizophrenia (SZ), major depressive disorder (MDD), attention-deficit/hyperactivity disorder (ADHD), and substance use disorders^20–24^. These associations highlight the critical importance of understanding the fundamental mechanisms of dopamine receptor signaling. In particular, biased signaling offers a promising framework for the development of targeted and effective therapeutics with fewer side effects, by selectively modulating disease-relevant signaling cascades^25^. Achieving this goal requires a deeper understanding of the molecular details related to how each D2-like receptor engages with different intracellular transducers.

The D2R is the most extensively studied subtype, with detailed characterization of its coupling to G_i/o/z_ proteins^16^, including differences between its two long and short splicing isoforms^19^. In contrast, D3R and D4R, despite sharing structural and signaling features with D2R, are less understood in terms of their selective coupling to Gα protein subtypes. In particular, the Gα protein coupling profile of D3R is debated, with some studies suggesting it fails to engage Gα_i_ subtypes and others reporting broader coupling across the G_i/o/z_ family^26–29^. In this study, we conclusively defined the signaling properties of D3R and D4R, demonstrating that D3R displays a distinct and consistent inability to couple with Gα_i_ proteins, despite robust activation of Gα_o_ and Gα_z_. This biased coupling pattern was further corroborated by exploring structural elements and defining the responsible molecular determinants. Finally, we demonstrated the physiological relevance of this discovery in neurons. Offering physiologically relevant evidence that D3R selectively couples to Gα_o_ and Gα_z_, we resolved previous inconsistencies and paved the way for the development of targeted D3R-based therapies.

## RESULTS

### Unique coupling profile of the dopamine D3 receptor

We initially assessed the G protein coupling specificity of the dopamine D2-like receptors (D2R, D3R, and D4R) to confirm their preferential interaction with heterotrimeric G proteins of the G_i/o/z_ subfamily, as previously reported^4^. To this goal, we transfected HEK293T cells with a representative Gα subunit from the four major G protein families, Gα_i/o/z_, Gα_s_, Gα_q_, and Gα_12/13_, and we applied a kinetic G protein nanoBRET assay to assess each G protein activation by D2R, D3R, and D4R in real time. Upon stimulation with either quinpirole (**Figure 1A**), a selective D2-like receptor agonist, or the non-selective endogenous agonist dopamine (**Supplementary** Figure 1A), all three receptors displayed clear and exclusive activation of the Gα_o_ subunit. No activation of Gα_s_, Gα_q_, or Gα_13_ proteins was observed under any condition, confirming the well-established Gα_i/o/z_-selective signaling profile of D2-like receptors. To further investigate the receptor-specific coupling preferences, we assessed the ability of D2R, D3R, and D4R to activate each of the six main Gα_i/o/z_ subunits (Gα_oA_, Gα_oB_, Gα_i1_, Gα_i2_, Gα_i3_, and Gα_z_). Applying the same G protein nanoBRET assay, we obtained concentration–response curves to quantitatively compare the coupling efficiency of the three dopamine receptors in response to both quinpirole and dopamine (**Figure 1B** and **Supplementary** Figure 1B). As expected, D2R and D4R displayed robust activation of all six G proteins with a preference for G_o_ over G_i_ and G_z_. In contrast, D3R showed a distinct and restricted coupling profile, with no detectable activation of any Gα_i_ subtype (Gα_i1_, Gα_i2_, and Gα_i3_), while retaining efficient coupling to Gα_oA_, Gα_oB_, and G_αz_ (**Figure 1B** and **Supplementary** Figure 1B). This allowed the calculation of EC_50_ and E_max_ values for individual G proteins, which are visualized as radar plots to facilitate comparison of both potency and efficacy (**Figure 1C**). These analyses further reinforced the functional similarity between D2R and D4R, while once again confirming the markedly different and selective profile of D3R. Taken together, these results suggest that D3R exhibits a selective signaling bias within the Gα_i/o/z_ family, favoring Gα_o/z_ over Gα_i_ subtypes.

**Figure 1.**
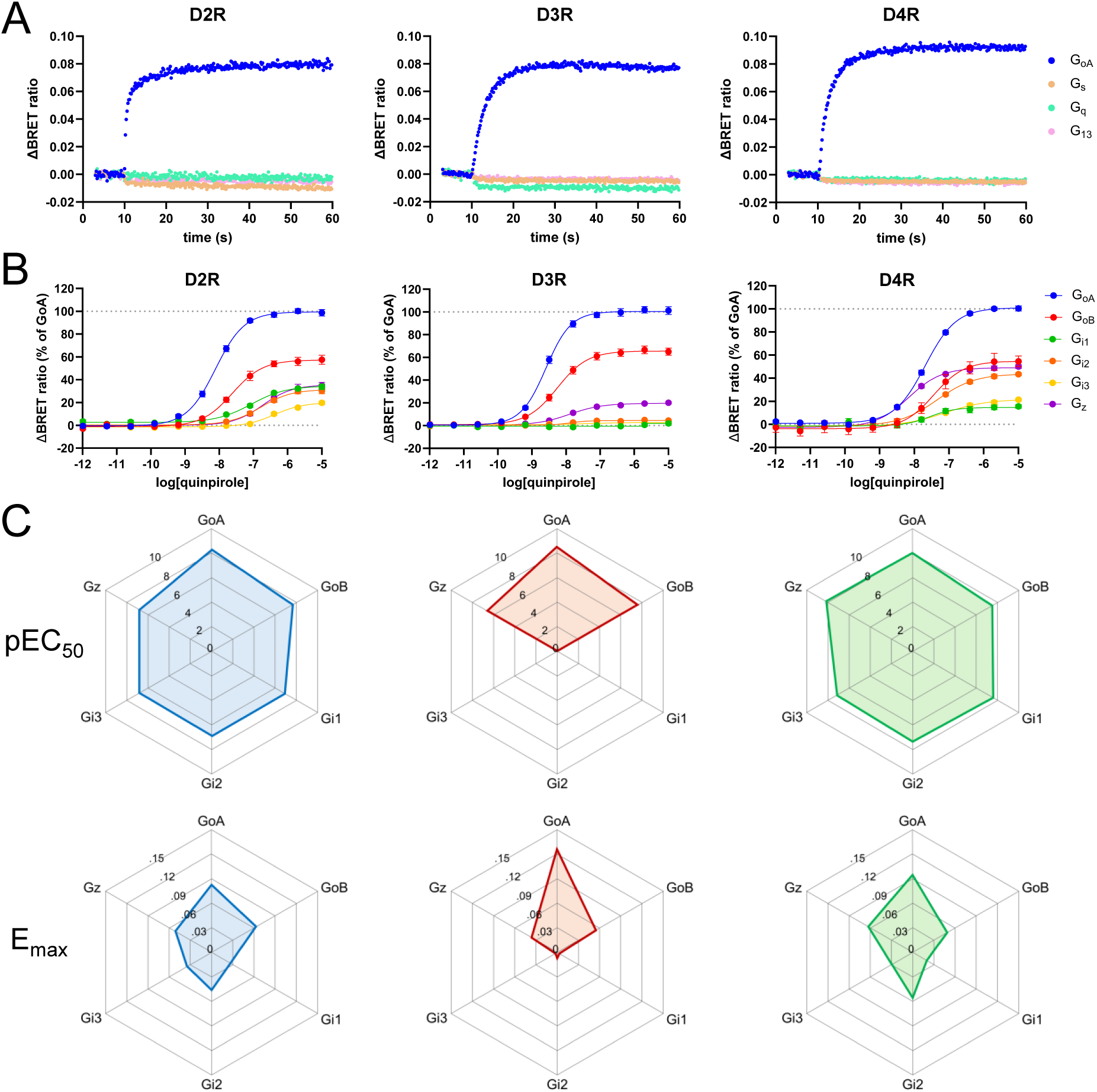
D3R shows a Gα_o/z_-selective coupling profile. (**A**) Representative kinetic activation profiles measured using the G protein nanoBRET assay for dopamine receptors D2R, D3R, and D4R in response to application of 10 µM quinpirole applied at 10 seconds. Co-transfected Gα subunits from the four major G protein families are indicated. (**B**) Concentration-response curves for D2R, D3R, and D4R co-transfected with indicated Gα_i/o/z_ subunits and stimulated with increasing concentrations of quinpirole. Data were normalized to the signal obtained with Gα_oA_ and shown as mean ± SEM. N=5 independent replicates. (**C**) Radar plots summarizing pEC_50_ and E_max_ for D2R, D3R, and D4R in response to quinpirole across all six Gα_i/o/z_ subunits.

The observation that D3R fails to couple to Gα_i_ subtypes, prompted us to investigate whether this reflects true molecular selectivity or results from suboptimal expression of either the receptor or the G protein in our system. We first tested whether varying the expression of D3R would rescue the coupling to Gα_i_. From this point onwards we used Gα_i2_ as a model to test D3R coupling. Accordingly, we transfected HEK293T cells with a fixed amount of either Gα_oA_ or Gα_i2_ plasmids and co-transfected increasing amounts of the D3R plasmid. We found that D3R robustly activated Gα_oA_ at all receptor expression levels tested, demonstrating the functionality of the assay across different transfection ratios (**Supplementary** Figure 2A). In stark contrast, no activation of Gα_i2_ was observed with any amount of transfected D3R-encoding plasmid (**Supplementary** Figure 2B). This suggests that receptor expression levels do not impact the assay ability to detect G protein activation effectively. To test whether the amount of Gα_i2_ was a limiting factor, we obtained concentration-response curves by treating cells transfected with a fixed amount of D3R and systematically increasing the levels of Gα_oA_, Gα_i2_, and Gα_z_. As a crucial negative control, no signal was generated in cells not transfected with Gα proteins. As expected, increasing the expression of Gα_oA_ and Gα_z_ resulted in a concentration-dependent decrease in the maximal response (E_max_) due to a shift of the equilibrium in the formation of the Gβγ-venus:GRK3-Nluc complex towards the G protein heterotrimer (**Supplementary** Figure 2C). In contrast, Gα_i2_ failed to produce any measurable activation signal, demonstrating that the lack of coupling is an intrinsic property and not an assay artifact. Taken together, these control experiments further substantiate that the observed Gα_o/z_-selective coupling profile is an intrinsic property of D3R.

### Ligand-bias does not explain the lack of Gα_i_ coupling of D3R

The concept of biased agonism proposes that ligands can stabilize distinct active receptor conformations, promoting selective engagement with specific transducers and activation of particular signaling pathways^12^. To determine whether quinpirole might act as a biased ligand by stabilizing D3R conformations incapable of coupling to Gα_i_ proteins, we compared its activity with multiple other dopamine receptor agonists. In addition to quinpirole and the endogenous ligand dopamine, we also included rotigotine, a potent D1R, D2R, and D3R agonist, which was recently used to induce an active conformation of D3R subsequently determined by cryo-EM structure^30,31^, sumanirole, a reported D2R-selective agonist, and ML417, a D3R-selective agonist. Using the G protein nanoBRET assay, we generated concentration-response curves for each ligand based on their ability to activate Gα_oA_, Gα_i2_, or Gα_z_ via D2R (**Figure 2A**) and D3R (**Figure 2B**). Notably, none of the tested agonists induced dissociation of the G_i_ heterotrimer in D3R-expressing cells, although all were capable of activating G_oA_ and G_z_ (**Figure 2B**). We further confirmed that ML417 preferentially activates D3R over D2R. While a partial activation of Gα_oA_ via D2R was observed at high ML417 concentrations, the ligand was 63-fold more potent at D3R and showed no activation of G_i2_ or G_z_. Additionally, we found that sumanirole triggered G_z_ heterotrimer dissociation with similar potency at both receptors but was five times more effective at activating G_oA_ via D2R compared to D3R. These findings exclude ligand bias as the cause of the inability of D3R to activate G_i_-dependent signaling.

**Figure 2.**
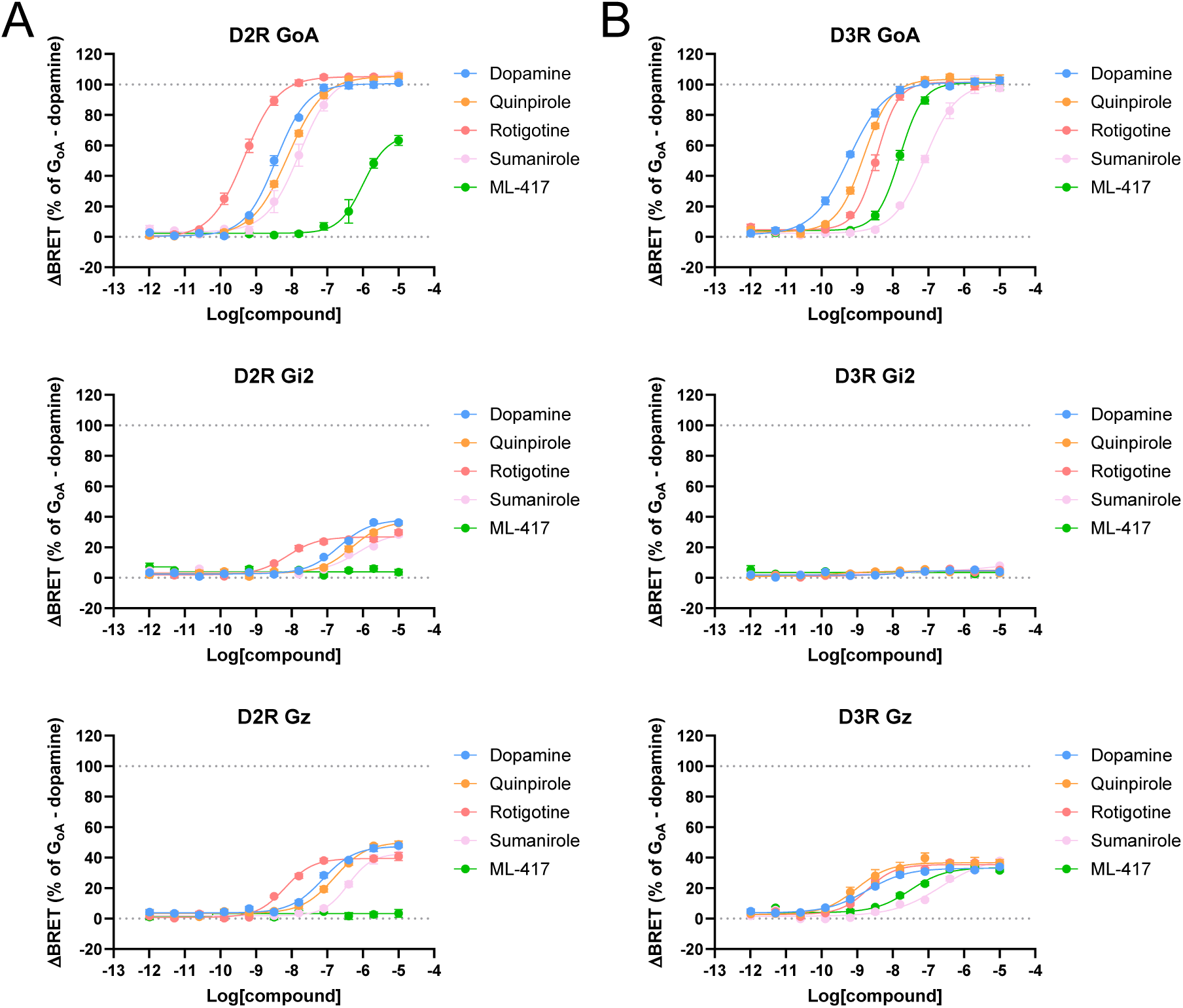
:The unique D3R activation profile is not a result of biased agonism. (**A**) Concentration-response curves obtained using the G protein nanoBRET assay in wild-type HEK293 cells expressing D2R with indicated G proteins and treated with indicated agonists. Data are shown as means ± SEM. N=5 independent replicates. (**B**) Concentration-response curves for D3R-transfected cells with indicated G proteins and treated with indicated agonists. Data were normalized to the signal obtained by applying dopamine to cells expressing Gα_oA_ and shown as mean ± SEM. N=5 independent replicates.

### Orthogonal GloSensor assay confirms lack of D3R-Gα_i2_ coupling

To further validate our G protein nanoBRET data, we employed the GloSensor cAMP inhibition assay in ΔGα_i/o/z_ cells to eliminate background interference from endogenous Gα_i/o/z_ proteins. Consistent with the nanoBRET data, stimulation with 1 µM quinpirole led to a marked decrease in cAMP levels across all the receptor-G protein combinations, except when D3R was co-expressed with Gα_i2_ (**Figure 3A–C**). The absence of cAMP inhibition in this condition further confirms that D3R does not functionally couple to Gα_i2_, supporting the conclusion that D3R preferentially signals through Gα_o_ and Gα_z_, rather than Gα_i_ subunits.

**Figure 3.**
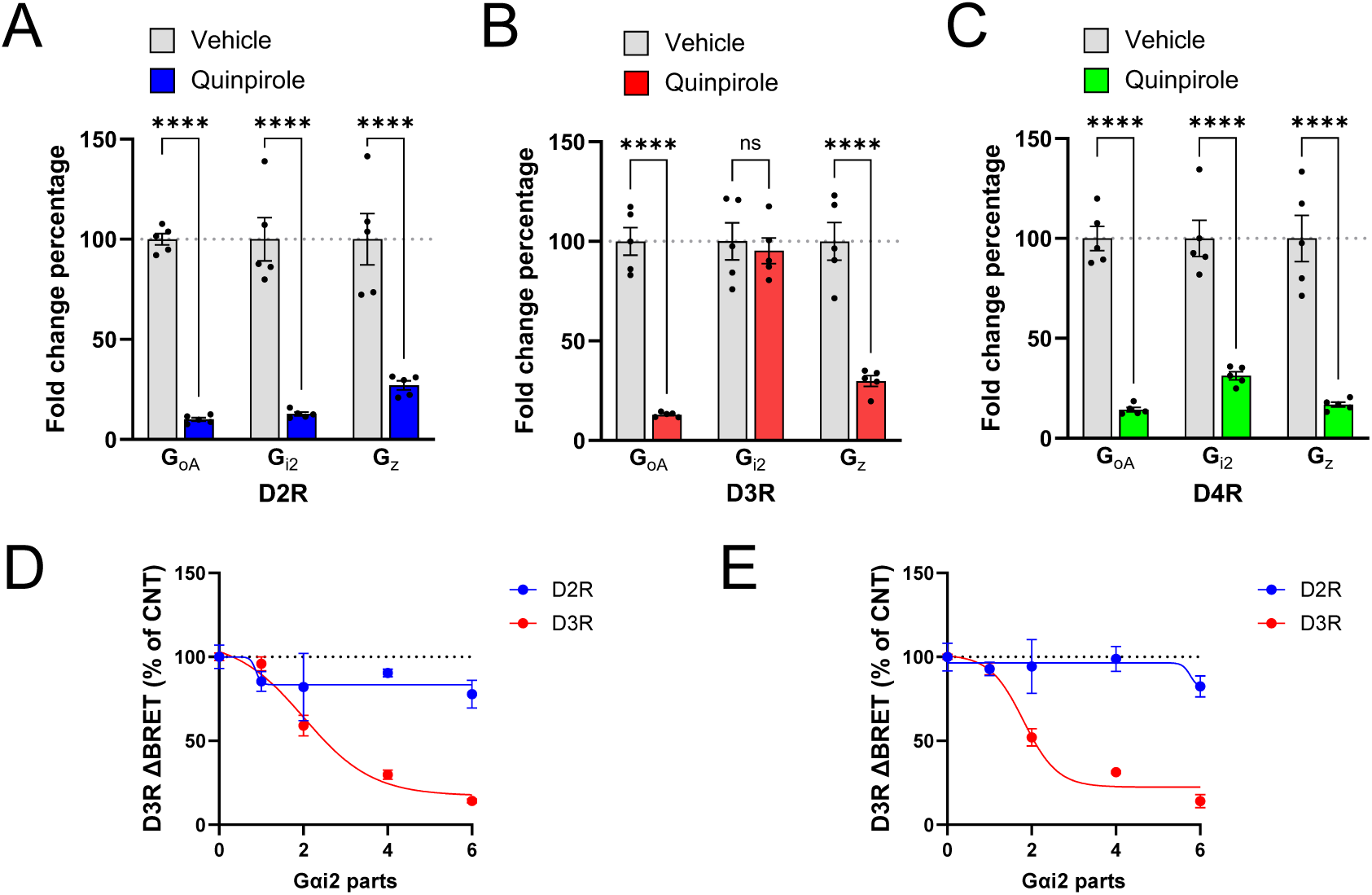
Orthogonal assays in ΔGα_i/o/z_ cells confirm lack of Gα_i2_ coupling by D3R. Percentage fold change of cAMP accumulation in response to stimulation with 1 µM quinpirole of D2R (**A**), D3R (**B**), D4R (**C**) co-transfected with G_oA_, G_i2_, and G_z_ in ΔGα_i/o/z_ HEK293 cells. Fold change was calculated as the ratio of maximal amplitude over baseline. cAMP levels were measured using the GloSensor assay in HEK293 cells lacking endogenous Gα_i/o/z_ proteins (ΔGα_i/o/z_ cells). Data are shown as mean ± SEM. N=5 independent replicates. ****P < 0.00001. (**D-E**) ΔGα_i/o/z_ HEK293 cells were co-transfected with either D2R or D3R, one part of a plasmid encoding for Gα_oA_ (208 ng) and increasing amounts of Gα_i2_ (208 ng for each part). G protein activation was measured using the G protein nanoBRET assay following stimulation with 1 µM quinpirole (**D**), or 1 µM dopamine (**E**). Data were normalized to 100% of the signal obtained by transfecting only Gα_oA_ (CNT). Data are shown as mean ± SEM. N=3 independent replicates.

To further confirm the selective coupling profile of D3R, we performed a competition experiment in ΔGα_i/o/z_ HEK293 cells co-transfected with either D2R or D3R, a constant amount of Gα_oA_, and increasing amounts of Gα_i2_. This setup divides the Gβγ-Venus sensor into two distinct pools: one associated with Gα_oA_ and the other with Gα_i2_. D2R, which couples to both G proteins, served as a control. As expected, increasing Gα_i2_ expression did not diminish D2R-mediated signaling in response to quinpirole (**Figure 3D**) or dopamine (**Figure 3E**), as D2R can activate both Gα pools, maintaining a strong overall signal. In contrast, D3R showed a markedly different profile. Increasing Gα_i2_ levels led to a concentration-dependent reduction in signal for both agonists, with over 86% reduction at the highest Gα_i2_ dose. This inhibition reflects the strict selectivity of D3R: because D3R cannot activate Gα_i2_, the excess Gα_i2_ sequesters Gβγ-Venus into inactive complexes, thereby limiting the Gβγ available for Gα_oA_-mediated signaling. These results further demonstrate that D3R does not functionally couple to Gα_i2_, even in the presence of excess Gα_i2_ protein.

### Structural determinants of D3R G protein coupling selectivity

Building on our previous findings, we sought to investigate the molecular determinants of the D3R-G protein coupling selectivity. We focused on the C-terminal α5 helix of the Gα subunit, a region known to be a primary interface for GPCR interaction^32–34^. An alignment of the Gα_oA_ and Gα_i2_ C-termini revealed key differences within the final 10 amino acids (**Figure 4A**). To test the importance of this region, we engineered two chimeric proteins by replacing the last 6 or 10 amino acids of the C-terminus of Gα_i2_ with the corresponding sequence from Gα_oA_, creating Gα_i2_Gα_oA(6)_ and Gα_i2_Gα_oA(10)_ (**Figure 4A-B**). To confirm the functional integrity of these chimeras, we assessed their activation by D2R in response to quinpirole (**Figure 4C**). As anticipated, both chimeras produced a clear activation signal, confirming productive coupling to the receptor. When tested with D3R, both chimeras exhibited a partial yet significant rescue of the coupling (**Figure 4D**). Specifically, Gα_i2_Gα_oA(6)_ restored 31% of the activation signal observed with wild-type Gα_oA_, while Gα_i2_Gα_oA(10)_ restored 37% of the signal. These results demonstrated that the Gα C-terminus is a critical determinant of coupling specificity for D3R. However, the incomplete nature of the rescue indicates that while these residues are essential, they are not the sole factor governing the interaction, suggesting that other regions of the G protein and possibly other contributors also influence the selective recognition.

**Figure 4.**
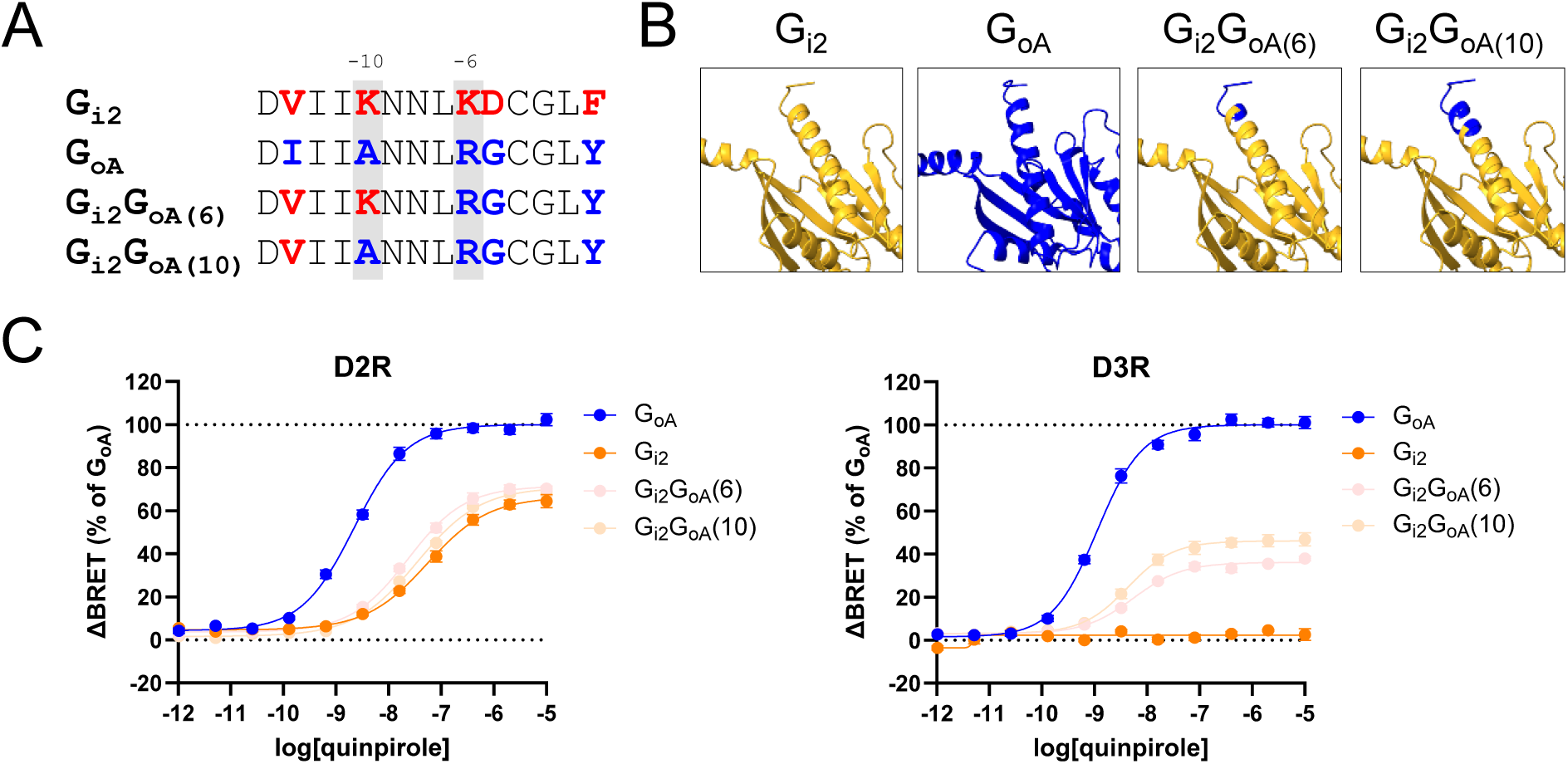
The C-terminal α5 helix (H5) is important for G protein coupling to dopamine receptors, but it is not the sole driver. (**A**) Alignment of the C-terminal α helix of Gα_i2_, Gα_oA_, and two chimeric constructs in which the last 6 or 10 residues of Gα_i2_ were replaced with the corresponding residues from Gα_oA_, respectively indicated as G_i2_G_oA_(6) and G_i2_G_oA_(10). Gα_i2_-specific residues are highlighted in red; GαoA-specific residues in blue. (**B**) Models of chimeric G protein constructs. (**C**) Concentration– response curves measured using the G protein nanoBRET assay for D2R (left) and D3R (right) co-transfected with Gα_oA_, Gα_i2_, G_i2_G_oA_(6), or G_i2_G_oA_(10). All experiments were conducted in wild-type HEK293T cells stimulated with increasing concentrations of quinpirole. Data were normalized to the signal obtained with Gα_oA_ and shown as mean ± SEM. N=5 independent replicates.

Having established the importance of the Gα C-terminus, we applied a similar approach to investigate the molecular determinants of the coupling selectivity of D3R. The intracellular loop 2 (ICL2) and ICL3 are known to be critical G protein interaction sites^35–38^. We therefore aligned the amino acid sequences of the ICL2 and ICL3 regions of D2R and D3R to identify differences and similarities (**Figure 5A**). Based on this analysis, we engineered six chimeric receptors by swapping ICL2 and/or ICL3 between D2R and D3R (**Figure 5B**). To confirm that these receptor chimeras retained the ability to respond to quinpirole and activate G proteins, we adopted the G protein nanoBRET and used G_oA_ as positive control. All six chimeric receptors produced a clear BRET signal upon stimulation, confirming they were properly expressed and functional (**Figure 5C**). The magnitude of activation was largely consistent with that of the wild-type receptor, except for the D3R^D2-ICL3^ chimera that showed significantly reduced coupling to Gα_oA_ (**Figure 5C**). Although further structural modeling is needed, we speculate that the reduced signaling may result from steric clashes, as the D3R^D2-ICL3^ chimera combines the longer ICL2 of D3R with the longer ICL3 of D2R. Nonetheless, this control confirmed that all chimeric receptors were functionally competent. We then co-transfected the wild-type and chimeric receptors with Gα_i2_ and analyzed their response to quinpirole (**Figure 5D**). The results revealed a striking pattern that depended exclusively on the origin of the ICL2. Every transfected receptor containing the ICL2 of D2R, such as wild-type D2R, D2R^D3-ICL3^, D3R^D2-ICL2^, and D3R^D2-ICL2/ICL3^, showed robust coupling to Gα_i2_. Conversely, every receptor containing the ICL2 of D3R, such as wild-type D3R, D3R^D2-ICL3^, D2R^D3-ICL2^, and D2R^D3-ICL2/ICL3^, showed no detectable Gα_i2_ activation. These findings pinpoint ICL2 as the primary molecular switch preventing the interaction between D3R and Gα_i_, thereby enforcing the selective signaling profile observed for D3R throughout this study.

**Figure 5.**
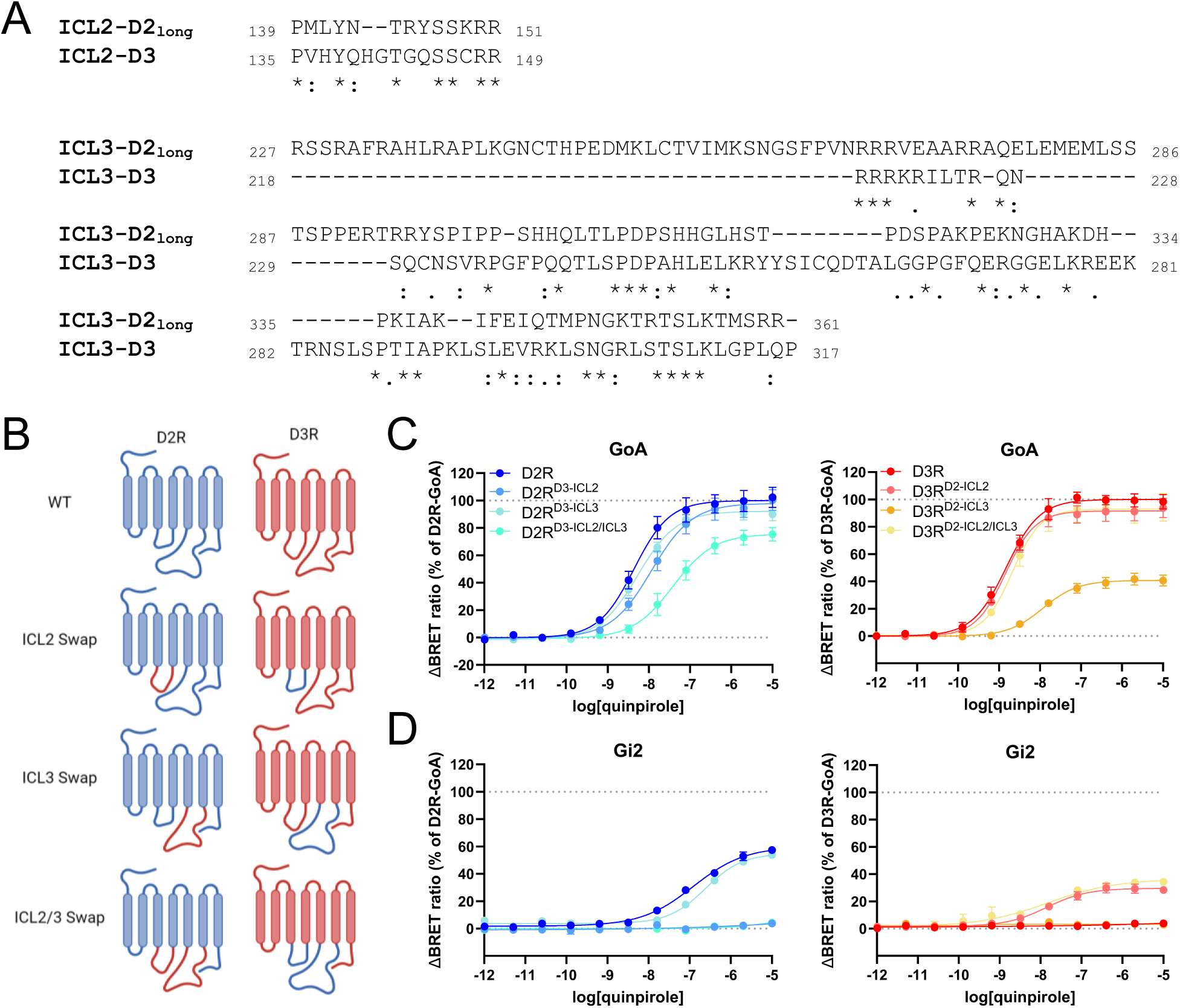
**Involvement of dopamine receptor intracellular loops in defining G protein selectivity.** (**A**) Sequence alignment of intracellular loop 2 (ICL2) and intracellular loop 3 (ICL3) of human D2R and D3R, highlighting the degree of conservation across the aligned sequences: *=full conservation, .=strong conservation, :=weak conservation. (**B**) Schematic representation of chimeric receptor constructs in which ICL2 and/or ICL3 of D2R and D3R were swapped. (**C**) Concentration–response curves obtained using the G protein nanoBRET assay for Gα_oA_ activation by wild-type and loop-swapped D2R and D3R constructs. (**D**) Corresponding concentration–response curves for Gα_i2_ activation. All experiments were conducted in wild-type HEK293T cells stimulated with increasing concentrations of quinpirole. Data were normalized to the signal obtained with Gα_oA_ and shown as mean ± SEM. N=5 independent replicates.

An investigation of the recently published structure of D3R bound to Gα_oA_-subtype of G protein (PDB:9F33) revealed that G350 is positioned in close proximity to a negatively charged cavity on D3R (**Figure 6A**). The presence of an Asp residue in this location in Gα_i_ might lead to electrostatic repulsion, destabilizing the interaction. In contrast, in D2R bound to Gα_i1_ (PDB: 7JVR), the corresponding Asp residue (D350) is stabilized by a surrounding positively charged cavity (**Figure 6A**). In addition to this, T142^ICL2^ on D3R is reported to get phosphorylated^39^, and the bulky phosphate group can be accommodated by a Gly residue. However, the presence of an Asp residue in this position would not only lead to electrostatic repulsion but also a steric clash. Interestingly, in the reported structures of D2R bound to either Gα_oA_ (PDB:8TZQ) or Gα_i1_ (PDB:7JVR), the corresponding T144^4.34^ exhibits a differential rotamer and points away from the Gα subunit (**Figure 6B**). Superimposing the structures reveals that A345, R349, and Y354 occupy a similar position in both D2R and D3R (**Figure 6C**) as well as the corresponding residues in Gα_i1_ in D3R-bound Gα_i1_ **(Figure 6C)**. While Y354 could potentially facilitate additional interactions with the surrounding positively charged pocket as compared to F354, the electron cloud over the benzene ring in both helps stabilize the G proteins in their corresponding receptor cavities.

**Figure 6.**
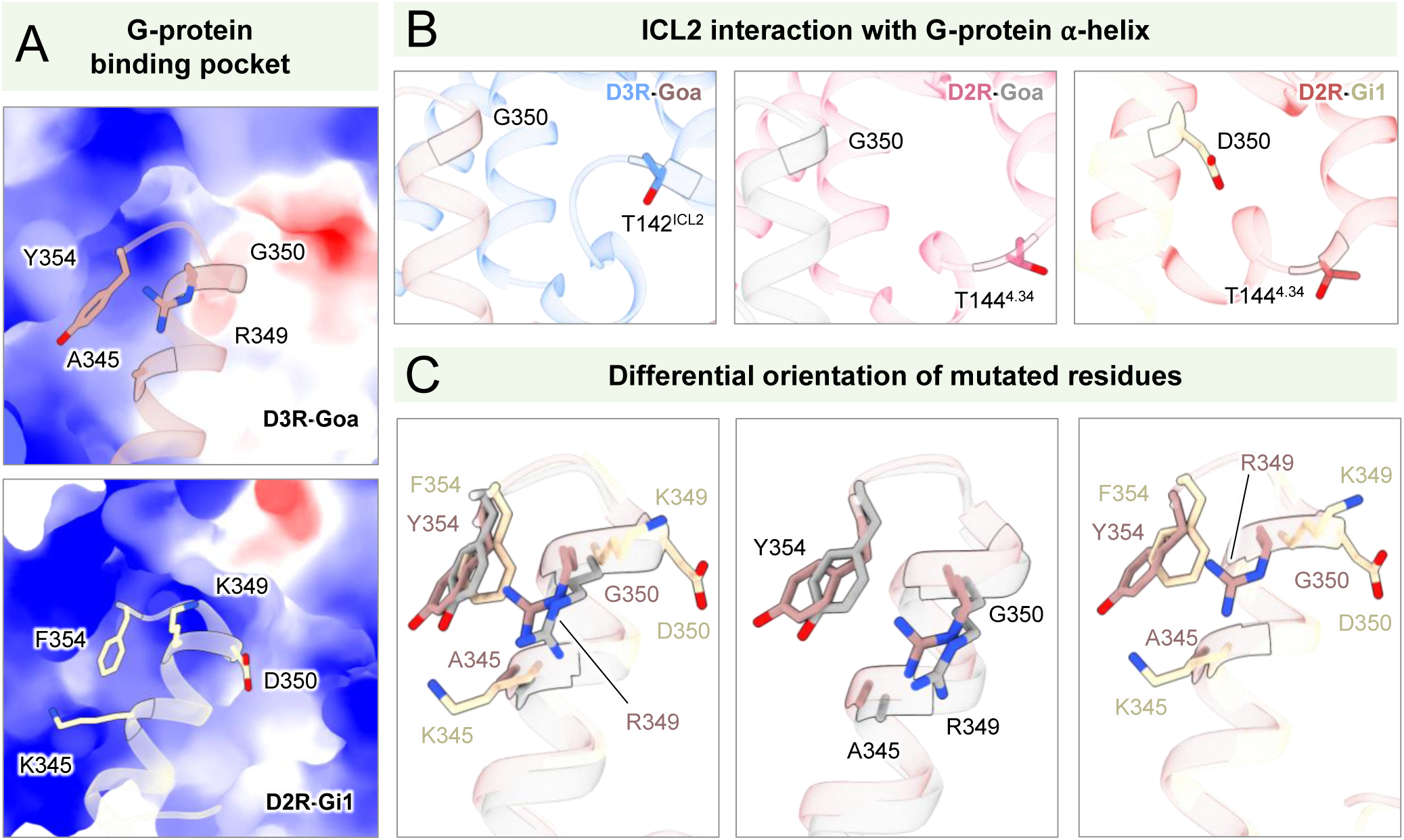
Structural comparison of the G protein binding cavity of D2R and D3R. (**A**) Surface representation of the charge distribution in the pocket occupied by Gα_oA_ in D3R (top) and Gα_i1_ in D2R (bottom). (**B**) T142^ICL2^ faces towards the ⍺5-helix in D2R, while T144^4.34^ in D3R exhibits a different rotamer, facing away from the ⍺5-helix. (**C**) Superimposition of the G⍺ subunit in the various structures reveals the orientation of the differing residues. (D3R-Gα_oA_: PDB – 9F33, D3R – cornflower blue, Gα_oA_ – rosy brown; D2R-Gα_oA_: PDB – 8TZQ, D2R – pale violet red, Gα_oA_ – gray; D2R-Gα_i1_: PDB – 7JVR, D2R – brick red, Gα_i1_ – peach puff).

### The unique coupling profiles of D2R and D3R produce distinct responses in hippocampal CA1 neurons

To examine the unexpected coupling profile of D3R in a physiological context, we focused on cAMP signaling in CA1 hippocampal neurons that richly express D2R and D3R^40^. We crossed the *cAMP Encoded Reporter* (*CAMPER*) mice, which conditionally express the ^T^EPac^VV^ FRET-based cAMP reporter^41^, with the CaMKIIα-Cre strain to achieve expression in the hippocampus^42^. Indeed, 2-photon imaging in acute brain slices from CaMKIIα-Cre:CAMPER^+/-^ revealed robust biosensor expression in the CA1 region (**Figure 7A**). We then used bath application of agonists at a low concentration (100 nM) to target D2R (sumanirole) or D3R (ML417) activation (**Figure 7B**). The Gα_s_-coupled β-adrenergic receptor was stimulated with isoproterenol (100 nM) as a control (**Figure 7B**). Sumanirole induced robust inhibition of cAMP, which is consistent with D2R-mediated Gα_i_ activation (**Figure 7C**). On the other hand, ML417 application paradoxically led to an increase in cAMP production (**Figure 7C**). As a control, the recording buffer alone exerted no change in cAMP dynamics. Subsequent application of isoproterenol increased cAMP in both the buffer and ML417 experiments, however the effect of isoproterenol was significantly greater after ML417 priming (**Figure 7C**). Pre-treatment with sumanirole completely blocked the effect of isoproterenol (**Figure 7C**). These data suggest sumanirole strongly inhibits AC activity via D2R-Gα_i_ and may display slow off-rate kinetics as cAMP inhibition persisted even in the presence of isoproterenol. The ability of ML417 to enhance cAMP supports the notion that D3R-Gα_o_ leads to release of Gβγ to conditionally sensitize AC activity^43^. This likely explains the sensitized cAMP response to isoproterenol following ML417 (**Figure 7D**). Collectively, these results capture how the exquisite coupling profiles of D2R and D3R contribute toward neuronal signal transduction.

**Figure 7.**
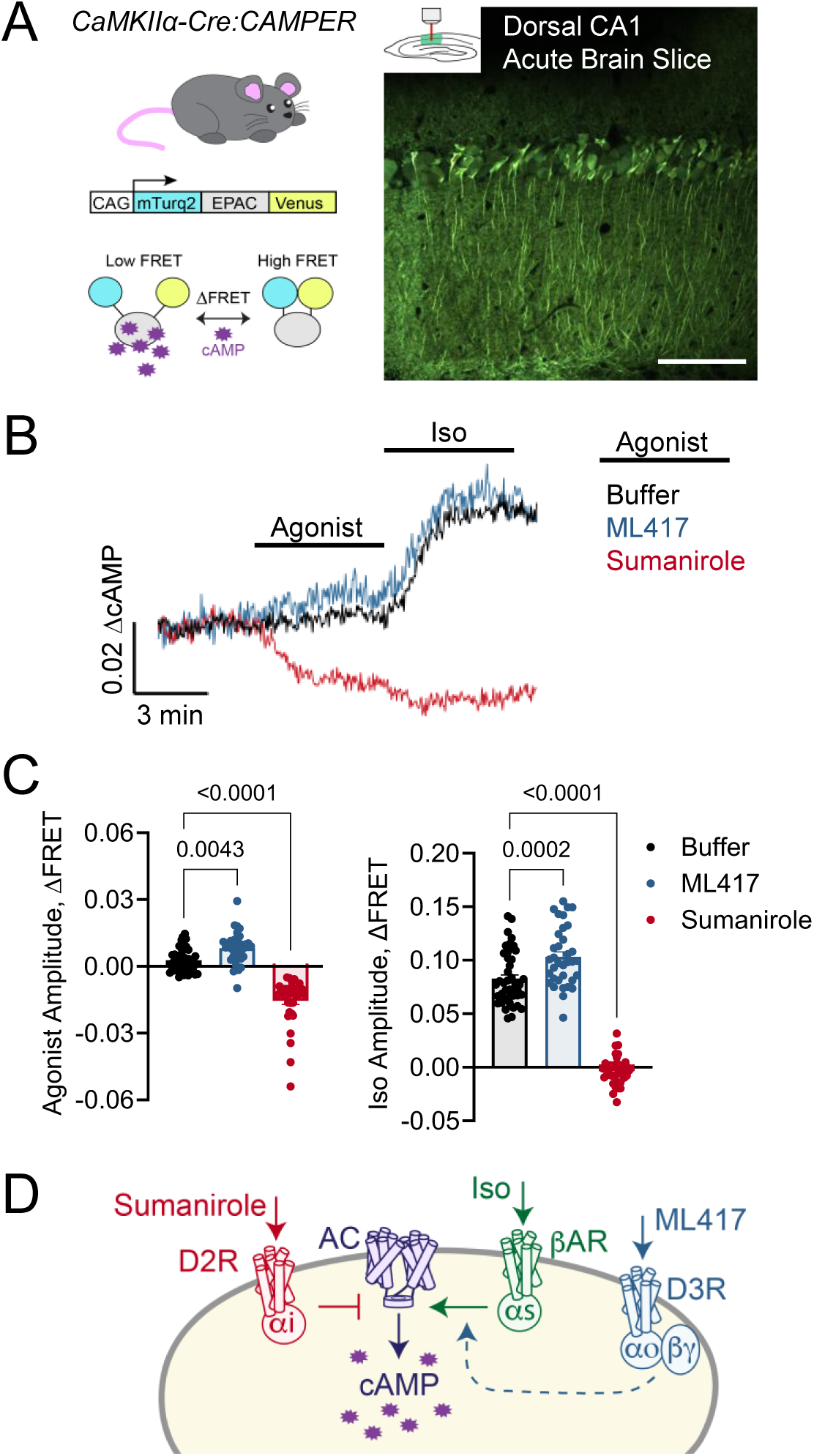
Agonism of D2R, but not D3R, inhibits cAMP signaling in CA1 hippocampal neurons. (A) Schematic of approach to visualize dorsal hippocampal CA1 neuronal cAMP dynamics in acute brain slices from CaMKIIα-Cre:CAMPER^+/-^ adult mice. Scale bar represents 100 microns. (B) Average trace of cAMP responses to agonist stimulation with ML417 500 (100 nM; n=35 neurons/6 mice), Sumanirole maleate (100 nM; n=38 neurons/6 mice), or control buffer (n=35 neurons/6 mice). In the same experiment slices were then immediately stimulated with Isoproterenol (Iso; 100 nM). (**C**) *left)* Maximum agonist induced cAMP amplitude calculated as peak agonist response minus basal cAMP level. One-way ANOVA was performed with Dunnett’s multiple comparisons test to buffer control. F=99.23, R^2^=0.6331, F (DFn, DFd)=1.365 (2, 115). *right)* Maximum Iso induced cAMP amplitude calculated as peak iso response minus peak agonist response. One-way ANOVA was performed with Dunnett’s multiple comparisons test to buffer control. F=239.4, R^2^=0.8063, F (DFn, DFd)=7.619 (2, 115). Data shown as mean ± SEM where each dot represents an individual neuron. Exact P values are depicted on the graphs. (**D**) Schematic illustration of D2R and D3R signal decoding in dorsal hippocampal CA1 neurons.

## DISCUSSION

Our study conclusively demonstrated that D3R is incapable of productive coupling with Gα protein subtypes Gα_i1_, Gα_i2_, and Gα_i3_, while it effectively couples to and activates Gα_o_ and Gα_z_. Although previous studies have partially addressed the G protein coupling selectivity of D3R, they produced inconsistent findings across different assay platforms, likely because of using modified systems that did not employ wild-type G proteins^26–28,31^. In detail, results from Lane and colleagues suggested a preferential coupling of D3R with Gα_oA_, minimal or absent activation of Gα_i1-3_ subtypes, while coupling to Gα_z_ was not assessed^26^. Their conclusions were supported by two sets of experiments. In the first, they engineered the tetracycline-inducible expression of PTX-resistant Gα subunits in HEK293 cells constitutively expressing D2R or D3R. Cells were treated with 10 µM dopamine, and receptor activity was assessed with a [^35^S]GTPγS binding assay. While their data suggested D3R does not couple to Gα_i2_, the induction of Gα_i2_ expression also elevated basal activity in unstimulated cells, complicating the interpretation of the results. Additionally, the engineered G proteins carried a point mutation in the α5 helix, a region critical for receptor interaction, raising concerns that the observed lack of Gα_i_ coupling could have been the reflection of using an artificial system. Using an alternative approach, they generated fusion proteins of dopamine receptors and PTX-insensitive G proteins to gain control over expression stoichiometry. However, elevated basal activity of D3R compared to D2R, combined with α5 helix modifications, again left unresolved the question of true Gα_i_ coupling. An independent group later applied miniG protein recruitment and G-CASE to profile the G protein coupling landscape of all five dopamine receptors^27^. Using a NanoBiT complementation assay based on miniG proteins, they detected miniG_o_ recruitment by D2R, D3R, and D4R, while miniG_i1_ was only recruited by D2R. However, application of the G-CASE assay resulted in discrepant observations of broader coupling, with D2R, D3R, and D4R all appearing to engage each member of the G_i/o/z_ family. Both assays have inherent limitations. MiniG proteins are truncated Gα subunits optimized for stable GPCR binding, but they lack the GTPase domain, which is a critical determinant of G protein coupling selectivity, thereby increasing promiscuity and reducing physiological relevance^44^. Similarly, the G-CASE assay does not employ wild-type G proteins, and Gα tagging positions may affect sensor performance, especially when the Gα:Gβγ interface undergoes subtle rearrangements rather than full dissociation^27^. Recent cryo-EM studies have resolved D3R-Gi complexes; however, such static structures can misrepresent physiological coupling, especially when stabilized using nanobodies, detergent micelles, or mutated receptor constructs^28,31^. Notably, signaling studies associated with these structures were obtained using the TRUPATH system, which shares similar design and limitations of the G-CASE assay, or with a NanoBiT recruitment performed in insect cells, which lack the mammalian post-translational modifications that could be crucial for authentic receptor behavior. In contrast, our study employed wild-type Gα proteins expressed in both parental and ΔGα_i/o/z_ HEK293 cells in two complementary functional assays. This experimental design, combined with the inclusion of multiple positive and negative controls, minimized artifacts associated with engineered or truncated Gα proteins and enabled a more physiologically relevant evaluation of receptor coupling selectivity. Our findings confirmed that D2R and D4R couple broadly to all Gα_i/o/z_ subtypes, while D3R exhibits exclusive coupling selectivity for Gα_oA_, Gα_oB_, and Gα_z_, with no detectable engagement with Gα_i1_, Gα_i2_, and Gα_i3_ under any tested condition.

Employing both G protein and receptor chimeras, we explored the molecular basis underlying this coupling selectivity. We confirmed the importance of the interface between the C-terminal α5 helix of the Gα proteins and the ICL2 of D3R in defining the biased coupling of D3R toward Gα_o_, while we showed that the ICL3 does not contribute to this selectivity. This is in conflict with previous findings that introducing 12 amino acids from the C-terminal segment of the ICL3 of D2R into the corresponding region of D3R partially restored Gα_i_ coupling^26^. Such discrepancies may arise from engineered features that bypass natural coupling constraints, highlighting the need for further molecular investigations. Models of the receptor-G protein interaction, based on available cryo-EM structures^28,31^, suggest that steric clashes and electrostatic repulsion may be responsible for the inability of D3R to engage Gα_i_ proteins. In particular, our analysis points at the involvement of D350 in Gα_i2_ and T142^ICL2^ in D3R, especially if phosphorylated by GRK2 as previously reported^39^. The role of these residues will require further investigations.

The Gα_o/z_-biased coupling of D3R has significant physiological implications. Although Gα subunits within the G_i/o/z_ family share high sequence similarity, they can engage distinct effectors and mediate non-redundant functions, as shown by the divergent phenotypes observed in Gα_o_, Gα_i_, and Gα_z_ knockout models and by recent studies characterizing unique intracellular effectors^43,45–50^. The restricted coupling profile of D3R suggests it may activate a distinct subset of signaling pathways limited to Gα_o/z_ and Gβγ-mediated responses, or it may bias signaling by excluding Gα_i_ subtypes, thereby enhancing the activation of alternative effectors, such as GIRK channels or Gβγ-sensitive adenylyl cyclases. As a promising target for treating substance use and other neuropsychiatric disorders^11,51,52^, D3R offers an additional layer of selectivity by modulating specific signaling pathways within defined cellular or subcellular contexts. Such enhanced selectivity holds the promise for the development of precisely targeted therapeutic strategies with reduced side effects. In this context, receptor-selective agonists are valuable both as research tools and potential treatments, offering improved outcomes compared to non-selective compounds due to differences in both receptor distribution and signaling^53^. Sumanirole, a D2R-selective agonist with 200-fold higher affinity for D2R over D3R^54^, exemplifies this approach by selectively initiating D2R-mediated effects in animal models, including prolactin release, regulation of striatal acetylcholine levels, and mediating autoreceptor-driven inhibition of nigrostriatal firing^54^. Similarly, pramipexole, a drug initially developed as a D2R agonist and later found to have high potency and selectivity for D3R^55^, has a demonstrated ability to delay L-DOPA-induced dyskinesia in the treatment of Parkinson’s disease^55–57^, suggesting that its clinical benefits may be partly due to D3R unique signaling properties. Although evidence for D3R-specific physiological effects is still emerging, the recent development of ML417 as a highly selective D3R agonist represents a major step forward in understanding D3R function and therapeutic potential^58^. ML417 has already been shown to possess neuroprotective properties, along with promising safety and pharmacokinetic profiles that would make it a strong lead compound for further drug development. Overall, D3R represents a unique case among GPCRs as no other GPCR with a primary coupling to G_i/o/z_ heterotrimeric proteins has been reported to be incapable of activating a subset of the family members. These insights not only clarify previous inconsistencies in the literature but also open new avenues for selective D3R pharmacology.

## METHODS

### Ethics statement

Procedures involving mice strictly followed NIH guidelines and were approved by the Institutional Animal Care and Use Committee at Augusta University. Mice were housed at consistent temperature with unlimited access to food and water under a constant 12-hour light/dark cycle. The following previously described strains were utilized: i.) *cAMP E*ncoded *R*eporter (*CAMPER*) (C57BL/6-Gt(ROSA)26Sortm1(CAG-ECFP*/Rapgef3/Venus*)Kama/J) (RRID:IMSR_JAX:032205)^41^, ii.) CaMKIIα-Cre (B6.Cg-Tg(Camk2a-cre)T29-1Stl/J) (RRID:IMSR_JAX:005359)^42^. The two strains were crossed to obtain experimental CaMKIIα-Cre:CAMPER^+/-^ mice. Both sexes were used and animal identification was verified by automated genotyping PCR (Transnetyx).

### DNA plasmids and chemicals

The plasmids encoding human codon-optimized DRD2, DRD3, and DRD4 were subcloned from the PRESTO-Tango library (Addgene; #66269, #66270, and #66271), a generous gift from Dr. Bryan Roth (UNC, Chapel Hill, NC). Gβ1-venus^156–239^ and Gγ2-venus^1-155^ were generous gifts from Dr. Nevin A. Lambert (Augusta University, Augusta, GA). Gα proteins and masGRK3CT-Nluc constructs were generous gifts from Dr. Kirill A. Martemyanov (UF Scripps Biomedical Research, Jupiter, FL). pGloSensor™-22F was purchased from Promega. Dopamine hydrochloride (#3548), quinpirole hydrochloride (#1061), and forskolin (#1099) were purchased from Tocris, while rotigotine (#HY-75502), sumanirole maleate (HY-70081A), and ML417 (HY-136390) were purchased from MedChemExpress. All chemicals were resuspended according to manufacturers’ instructions, aliquoted, and stored at -20°C until use.

### Cell cultures and transfections

HEK293T/17 cells were purchased from ATCC, while Gα_i/o/z_-deficient HEK293A cells (ΔGα_i/o/z_), a generous gift from Dr. Asuka Inoue (Tohoku University, Japan), were previously generated using three rounds of CRISPR–Cas9 mutagenesis and validated^59^. Cells were cultured in DMEM (Gibco, 10567-014) supplemented with 10% FBS (Biowest, S1520), non-essential amino acids (Gibco, 11140-050), 100 units/ml penicillin and 100 µg/ml streptomycin (Gibco, 15140-122), and 250 µg/ml amphotericin B (ThermoFisher, 15290-018) at 37°C and 5% CO_2_. For transfection, two million cells were seeded in each well of 6-well plates in 1.5 mL medium per well without antibiotics and containing 10% dialyzed FBS (Biowest, S181D) for 4 hours and then transfected with a mixture containing a 1:3 ratio of DNA plasmid (2.5 µg) and polyethylenimine (PEI; 7.5 µl) (VWR, AAA43896) in a final volume of 500 µL per well. The following amount of each plasmid was transfected, unless otherwise indicated: 0.21 µg GPCR (D2R, D3R, or D4R), 0.83 µg Gα protein (Gα_oA_, Gα_oB_, Gα_i1_, Gα_i2_, Gα_i3_, Gα_z,_ Gα_s_, Gα_q_, Gα_12_), 0.21 µg Gβ1-venus^156–239^, 0.21 µg Gγ2-venus^1-155^, and 0.013 µg masGRK3CT-Nluc. An empty vector, pcDNA3.1, was used to normalize the ratio of transfected plasmids. Transiently transfected cells were incubated for 18-22 hours before being tested.

### G protein nanoBRET assay

The day after transfection, cells were briefly washed with PBS, resuspended in BRET buffer (PBS supplemented with 0.5 mM MgCl_2_ and 0.1% glucose), collected in 1.5 ml tubes, and centrifuged for 5 minutes at 500 x g. Pelleted cells were resuspended in 300 µl of BRET buffer, and 25 µl of cells were plated in 96-well white microplates (Greiner Bio-One). The Nluc substrate Hikarazine-103 was purchased from Synthelis and used according to the manufacturer’s instructions. BRET measurements were obtained using a POLARstar Omega microplate reader (BMG Labtech). All measurements were performed at room temperature, and the BRET signal was determined by calculating the ratio of the light emitted by Gβ1γ2-venus (collected using the emission filter 535/30) to the light emitted by masGRK3CT-Nluc (475/30). In kinetics assays, the baseline value (basal BRET ratio) was averaged from recordings of the five seconds before agonist injection. In concentration-response experiments, 25 µl of cells per well were plated and mixed with the Nluc substrate. Initial readings were performed to establish basal BRET ratio, and then agonists were added. The BRET signal was recorded for 60 seconds. ΔBRET ratios were obtained by subtracting the basal BRET ratio from the maximal amplitude measured.

### cAMP inhibition assay

1 million ΔGα_i_ cells were seeded in each well of 6-well plates. 4 hours later, cells were transfected with plasmids for mammalian expression of individual dopamine receptors (D2R, D3R, or D4R), pGloSensor™-22F cAMP plasmid (Promega), and individual Gα proteins Gα_oA_, Gα_i2_, or Gα_z_. 18-22 hours after transfection, cells were collected in PBS, centrifuged at 500 x g for 5 minutes at room temperature, and the supernatant was discarded. The pelleted cells were resuspended in 300 μl of BRET buffer (PBS supplemented with 0.5 mM MgCl_2_ and 0.1% glucose), and 40 μl was plated into each well of a white 96-well plate, followed by the addition of 10 μl of D-Luciferin potassium salt substrate (GoldBio; #LUCK-100; final concentration 600 µg/ml). After incubation at 37°C for 1h, the plate was placed in a BMG Omega microplate reader and maintained at 28°C until a stable baseline value was recorded. Cells were then treated with 1 μM quinpirole for 30 seconds before adding 0.5 μM forskolin and recording luminescence for 25 minutes at 72-second intervals. A fold change (FC) was calculated for each sample by dividing the maximum relative light units (RLU) obtained after forskolin application by the average baseline RLU. For the quinpirole concentration-response curve, the FC for samples treated with the vehicle was averaged and considered 100% of forskolin-induced cAMP levels. The FC of all other samples was then normalized to the vehicle control FC average and expressed as a percentage of cAMP inhibition.

### 2-photon FRET cAMP imaging

Acute brain slices were prepared from adult mice (2-4 months of age) as similarly described^60,61^. Animals were isoflurane anesthetized followed by decapitation and rapid extraction of the brain, which was subsequently agarose-mounted on a vibratome (Precisionary VF-310-0Z) in ice-cold oxygenated buffer consisting of (in mM): KCl (2.5), NMDG (93), glucose (25), HEPES (20), sodium ascorbate (5), sodium pyruvate (3), thiourea (2), NaH_2_PO_4_ (1.2), CaCl_2_ (0.5), MgCl_2_ (10), NaHCO_3_ (30). 300-micron thick coronal slices containing the hippocampus were sectioned and incubated for 1 hour at 34°C in an oxygenated recovery buffer consisting of (in mM): NaCl (126), KCl (2.5), CaCl_2_ (2), MgCl_2_ (2), NaHCO_3_ (18), NaH_2_PO_4_ (1.2), glucose (10). Slices were then maintained at ambient temperature oxygenated recording buffer consisting of (in mM): NaCl (125), KCl (2.5), CaCl_2_ (2), MgCl_2_ (2), NaH_2_PO_4_ (1.25), NaHCO_3_ (25), glucose (25). Individual brain sections were then constantly perfused in recording buffer at approximately 2 ml per minute in a recording chamber (Warner Instruments) for FRET imaging on a Zeiss 780 multiphoton confocal microscope (20X W Plan-Apochromat objective). Excitation of the cAMP FRET donor (mTurquoise2) was achieved by tuning a Ti:Sapphire laser (Coherent Chameleon Vision S) to 850 nm. Individual photomultiplier tubes were utilized to simultaneously capture emission from the FRET donor (455-509 nm) and FRET acceptor (Venus; 526-571 nm) at 2.5 second intervals. ML417 (MedChemExpress HY-136390), Sumanirole maleate (MedChemExpress HY-70081A), and isoproterenol hydrochloride (Iso; TCI America I0260) were bath applied as indicated in the text. FRET values were calculated using standard ImageJ tools from the raw fluorophore intensity at the neuronal cell body. Non-responsive cells were excluded based on a previously established cutoff criterion (two multiplied by the standard deviation of the baseline prior to drug treatment)^62,63^.

### Statistical analysis

Statistical analysis was performed using GraphPad Prism version 10 software. Concentration-response curves were fitted to a sigmoidal four parameter logistic function (variable slope analysis) to quantify agonist potencies (pEC_50_), maximal responses (E_max_). Concentration-response curves were fitted and Hill slope values for each agonist were close to 1. At least three independent biological replicates were used for each experiment.

## Data Availability Statement

All processed data that support the findings of this study are available within the paper and its supplemental data. Any additional information required to reanalyze the data reported in this paper is available from the corresponding author upon request.

## Supporting information

Zanetti Supplementary Figures

## Acknowledgments

This work was supported by NIH awards R01 DC022104 to C.O and R01 NS129554 to B.S.M. We thank Dr. Asuka Inoue at Tohoku University for the CRISPR-knockout ΔGα_i/o/z_ HEK293 cell line under a Materials Transfer Agreement. The CAMPER mouse strain was a gift from Dr. Kirill Martemyanov. The Cell Imaging Core facility at Augusta University was utilized for 2P imaging.

