## Supplementary material for "Go/z-biased coupling profile of the dopamine D3 receptor": Zanetti Supplementary Figures

### Supplementary Figure 1

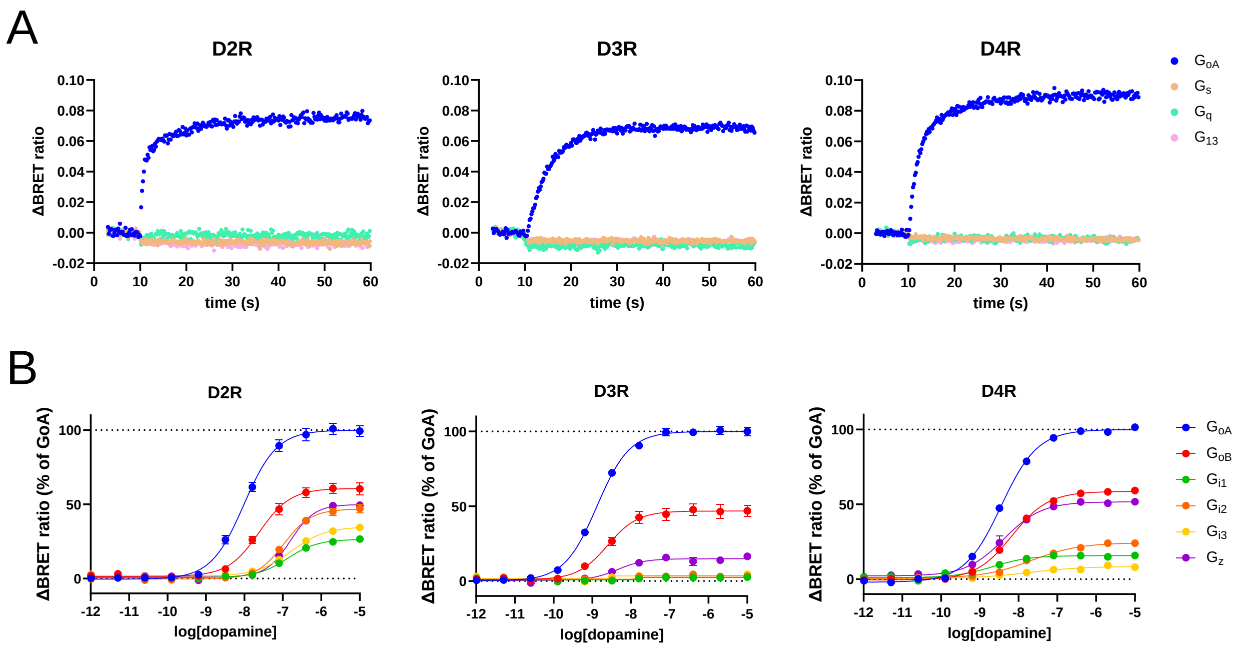


**Supplementary Figure 1. Dopamine-mediated activation of $G_{i/o/z}$ family members by D2R, D3R, and D4R.** (**A**) Kinetics curves of dopamine receptor activation of a representative heterotrimeric G protein for each family. 10 µM dopamine application at 10 seconds stimulates the activation of GoA but not Gs, Gq, or G13 by D2R, D3R and D4R. (**B**) Concentration-response curves showing D2R, D3R, and D4R activation of each $G_{i/o/z}$ family member by dopamine. Data are shown as mean ± SEM of values normalized to $G\alpha_{oA}$ signal. N=5 independent replicates.

### Supplementary Figure 2

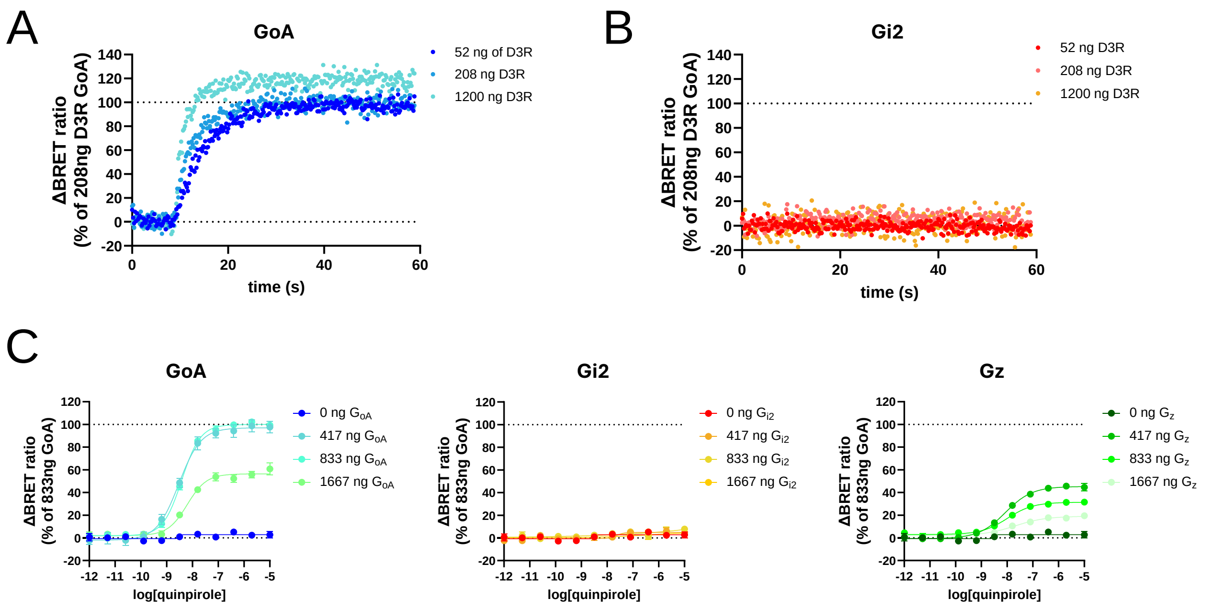


**Supplementary Figure 2. Lack of G$\alpha_{i2}$ activation by D3R is independent of receptor or G protein expression levels.** (**A-B**) To assess the impact of receptor expression levels, increasing amounts of DNA plasmid encoding D3R were transfected over 2500 ng of total DNA. Empty vector pcDNA3.1 was co-transfected to maintain a constant amount of total DNA transfected. Cells were stimulated with 100 µM dopamine at 10 seconds. Representative kinetic activation profiles measured using the G protein nanoBRET assay for D3R co-transfected with either G$\alpha_{oA}$ (**A**, positive control) or G$\alpha_{i2}$ (**B**). (**C**) Concentration–response curves measured using the G protein nanoBRET assay for a fixed amount of D3R co-transfected with increasing amounts of DNA plasmids encoding either G$\alpha_{oA}$, G$\alpha_{i2}$, or G$\alpha_z$. Empty vector pcDNA3.1 was co-transfected to maintain a constant amount of total DNA transfected. Cells were stimulated with increasing concentrations of quinpirole. Data were normalized to the signal obtained by transfecting 833 ng of G$\alpha_{oA}$ and shown as mean ± SEM. N=5 independent replicates.
